# Gut microbiota analyses of Saudi populations for type 2 diabetes-related phenotypes reveals significant association

**DOI:** 10.1101/2021.10.25.465666

**Authors:** Fahd A Al-Muhanna, Alexa K. Dodwell, Abdulmohsen H Al Eleq, Waleed I Albaker, Andrew W. Brooks, Ali I Al-Sultan, Abdullah M Al-Rubaish, Khaled R Alkharsah, Raed M Sulaiman, Abdulaziz A Al-Quorain, Cyril Cyrus, Rudaynah A Alali, Chittibabu Vatte, Fred L. Robinson, Carrie Nguyen, Xin Zhou, Michael P. Snyder, Afnan F Almuhanna, Brendan J Keating, Brian D. Piening, Amein K Al-Ali

## Abstract

Large-scale gut microbiome sequencing has revealed key links between microbiome dysfunction and metabolic diseases such as T2D. To date, these efforts have largely focused on Western populations, with few studies assessing T2D microbiota associations in Middle Eastern communities where T2D prevalence is now over 20%. We analyzed the composition of stool 16S rRNA from 461 T2D and 119 non-T2Dparticipants from the Eastern Province of Saudi Arabia. We quantified the abundance of microbial communities to examine any significant differences between subpopulations of samples based on diabetes status and glucose level. We observed overall positive enrichment within diabetics compared to healthy individuals and amongst diabetic participants; those with high glucose levels exhibited slightly more positive enrichment compared to those at lower risk of fasting hyperglycemia. In particular, the genus *Firmicutes* was upregulated in diabetic participants compared to non-diabetic participants, and T2D was associated with an elevated *Firmicutes/Bacteroidetes* ratio, consistent with previous findings. Based on diabetes status and glucose levels of Saudi participants, relatively stable differences in stool composition were perceived by differential abundance and alpha diversity measures.

**Author summary:** The rates of Type 2 diabetes (T2D) in Saudi Arabia have risen dramatically in the last several decades due to socio-economic changes resulting in changes in dietary and sedentary lifestyles. This emergence has grown more rapidly and affects larger proportions of the population with estimates of T2D prevalence impacting 25% of the population. There is a paucity of microbiome data from Middle Eastern populations, and previous studies have been conducted on small sample sizes. Here we report on the first-ever characterization of gut microbiota T2D versus non-T2D and largest microbiome study ever conducted in a Middle Eastern country. The datasets from this study are important to create a regional reference T2D-microbiome catalogue which will propel the understanding of regional gut flora which are associated with T2D development. Based on T2D status and quantified glucose levels of Middle Eastern participants, relatively stable differences in stool composition were observed by differential abundance and alpha diversity measures. Comparing overlapping and varying patterns in gut microbiota with other studies is critical to assessing novel treatment options in light of a rapidly growing T2D health epidemic.

## Introduction

The human gut hosts 100 trillion microorganisms, encompassing thousands of species collectively, weighing an average 1.5 kg per person [1,2]. The human microbiota is important because of its metagenomic repertoire, which is estimated to be 100 times larger than the human genome and encodes a vast array of functionality critical for host physiology and metabolism [2]. The bacterial components responsible for triggering theses physiological functions are currently the subject of intensive research. Differences in human gut microbiome composition have been linked to metabolic diseases such as T2D and obesity [3-7]. Identifying specific bacterial biomarkers within the microbiome could help predict the occurrence of T2D or tailor treatments in high-risk subjects to prevent or delay the onset of metabolic diseases. The molecular mechanisms through which the intestinal microbiota play a key role in metabolic diseases are linked to an increased energy harvesting and the triggering of the low-grade inflammatory status characterizing insulin resistance and obesity [8-9].

The prevalence of T2D is increasing worldwide, with current data indicating that at least 8.5% of the world’s population is affected, with the worldwide prevalence expected to reach 12% by 2025 [10-11]. T2D is mainly caused by insulin resistance and relative insulin deficiency [12]. Saudi Arabia, with a total population of over 20 million, has an estimated T2D constituting 25% of the total population [13]. The rapid rate of increase of T2D disease in some areas of Saudi Arabia, which increased from 16% in 2005 to over 25% in 2011, is thought to be due to rapid lifestyle changes such as diet and sedentary lifestyle, as we;; as adverse environmental factors [13].

We analyzed the composition of 16S rRNA from the stool samples collected from Saudi Arabian participants residing in the Eastern Province and quantified the abundance of microbial communities to determine significant differences between subpopulations of samples based on diabetes status and glucose level. We assessed alpha diversity between the subpopulations to measure species richness and evenness among samples noting that an increased *Firmicutes:Bacteriodetes* ratio has previously been observed in the microbiota of obese/diabetic individuals compared to the microbiota of healthy individuals. Furthermore, individuals with diabetes were tracked for high glucose level (>126 mg/dL) as it is an indicator of fasting hyperglycemia, which could potentially lead to severe long-term complications including cardiovascular disease, neuropathy and kidney failure.

## RESULTS

Principal coordinate analysis (PCoA) of the generated 16S datasets is shown in Fig S2. The first and second principal coordinated explained 25% and 7%; 29% and 7% and 34% and 6% of the Diabetes Status and Gender variance, respectively. Levels of the 150 most abundance microbial genera within T2D and non-T2D participants were observed to differ significantly in stool microbiota abundance derived from 16S sequencing (Figures S3a and S3b).

Fig 1 (a and b) shows the rank abundant curve and Permutational Multivariate Analysis of Variance (PERMANOVA) cloud, respectively for Saudi T2D and control 16S stool microbiota datasets. These show that the microbiome communities differ globally between T2D and non-T2D subjects at statistical significance, *p* = 0.01. The abundance of Taxonomic Composition in males and females is clearly evident in both females (Figures S4a and S4b) and in males (Figures S5a and S5b). We also compared Saudi T2D participants with higher glucose >126 mg/dL versus lower glucose strata <=126 mg/dL glucose using the top 150 genera. Amongst the 298 samples with glucose data, n=193 were in the higher glucose strata and n=105 were in the lower strata (Figure S6). Unlike previous studies conducted on Western populations, the Saudi participants with T2D and higher glucose levels showed a trend toward increased diversity, a result that is similar to another recently reported study from a United Arab Emirates (UAE) cohort [14].

**Figure 1:**
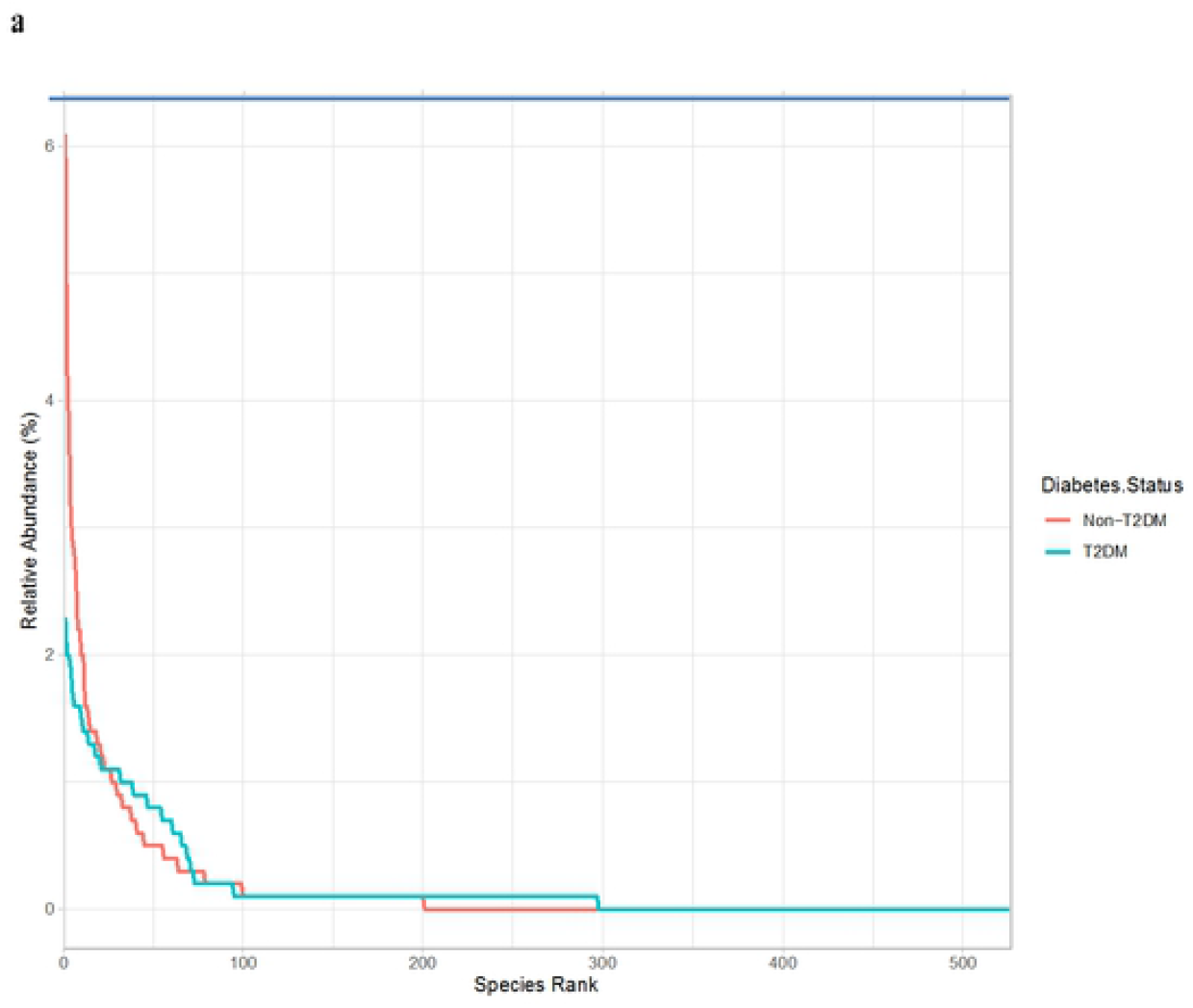
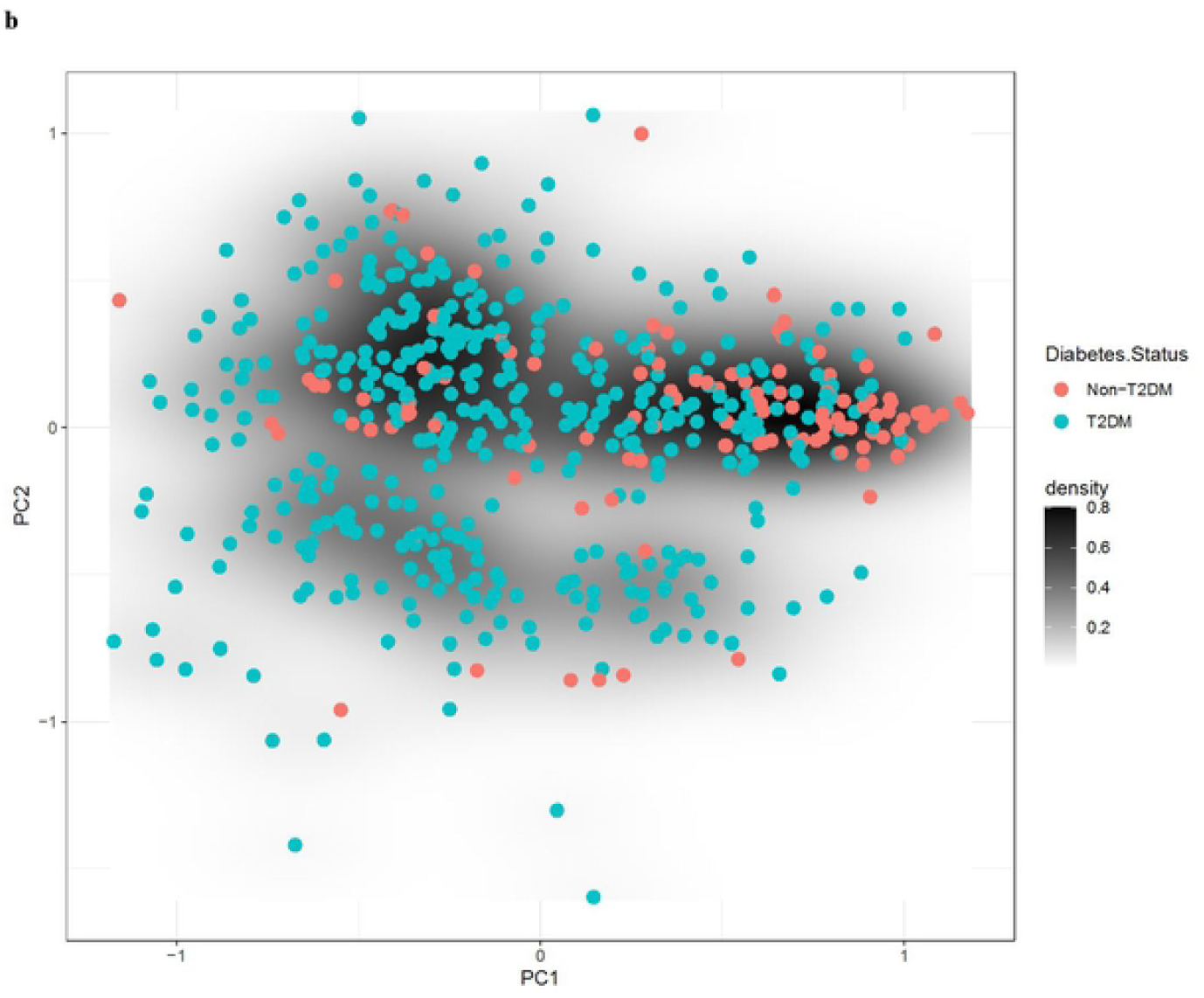
Rank abundant curve (a) and permutational multivariate analysis of variance cloud (b) for Saudi T2D and control 16S stool microbiota datasets. This figure shows the rank abundant curve and Permutational Multivariate Analysis of Variance (PERMANOVA) cloud respectively for Saudi T2D and control 16S stool microbiota datasets. These show that the microbiome communities differ globally between T2D and non-T2D subjects at statistical significance, *p* = 0.01.

Alpha diversity was compared in males versus females (n = 204 and 226, respectively) with no significant differences observed using various different classifications: ACE (Abundance-based Coverage Estimator) and Chao1 indices to estimate richness (measurement of OTUs expected in samples given all the bacterial species identified in the samples); Shannon-Weaver, Simpson and Inverse Simpson to define different levels of resolution (phylum, class, order, family, genus, and species); and Fisher (Fig S7). Alpha diversity of T2D versus non-T2D participants revealed statistically significant enrichment of the Shannon-Weaver and Simpson metrics (Figure S8) (p < 2.26 × 10^−10^ (CI: -0.392 to -0.718)) and p < 4.63 × 10^−7^ (CI: -0.049 to -0.108) for Shannon and Simpson diversity, respectively. Saudi T2D cases versus controls showed an association with an elevated *Bacteroidetes/Firmicutes* ratio, p = 2.2 × 10^−5^ t-test (Fig S9).

We observed an overall positive enrichment of microbiota genus/families for diabetics compared to healthy individuals. In addition, among T2D patients, those with high glucose levels exhibited slightly more positive enrichment compared to those at lower risk of fasting hyperglycemia (Fig 2a and 2b and Table S1). In particular, the *Akkermansia, Acidaminococcus, Megamonas, Dialister, Lactobacillus and Paraprevotella* genus were enriched at p < 1 × 10^−9^ in T2D versus non-T2D. The *Fusobacterium, Dialister, Akkermansia* and *Prevotella* genus were enriched in low versus high-risk T2D using a fasting glucose cutoff of 126 mg/dL.

**Figure 2:**
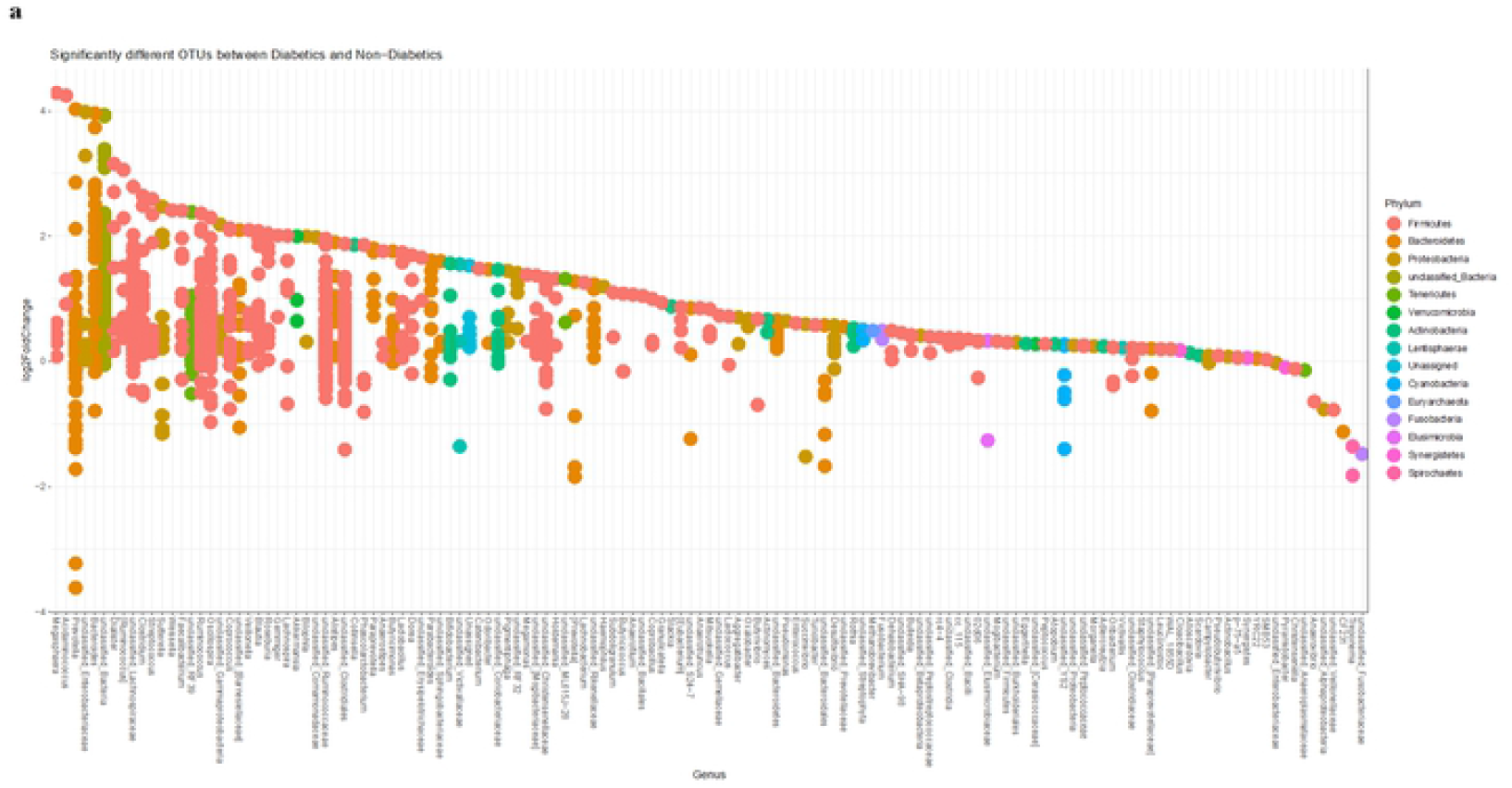
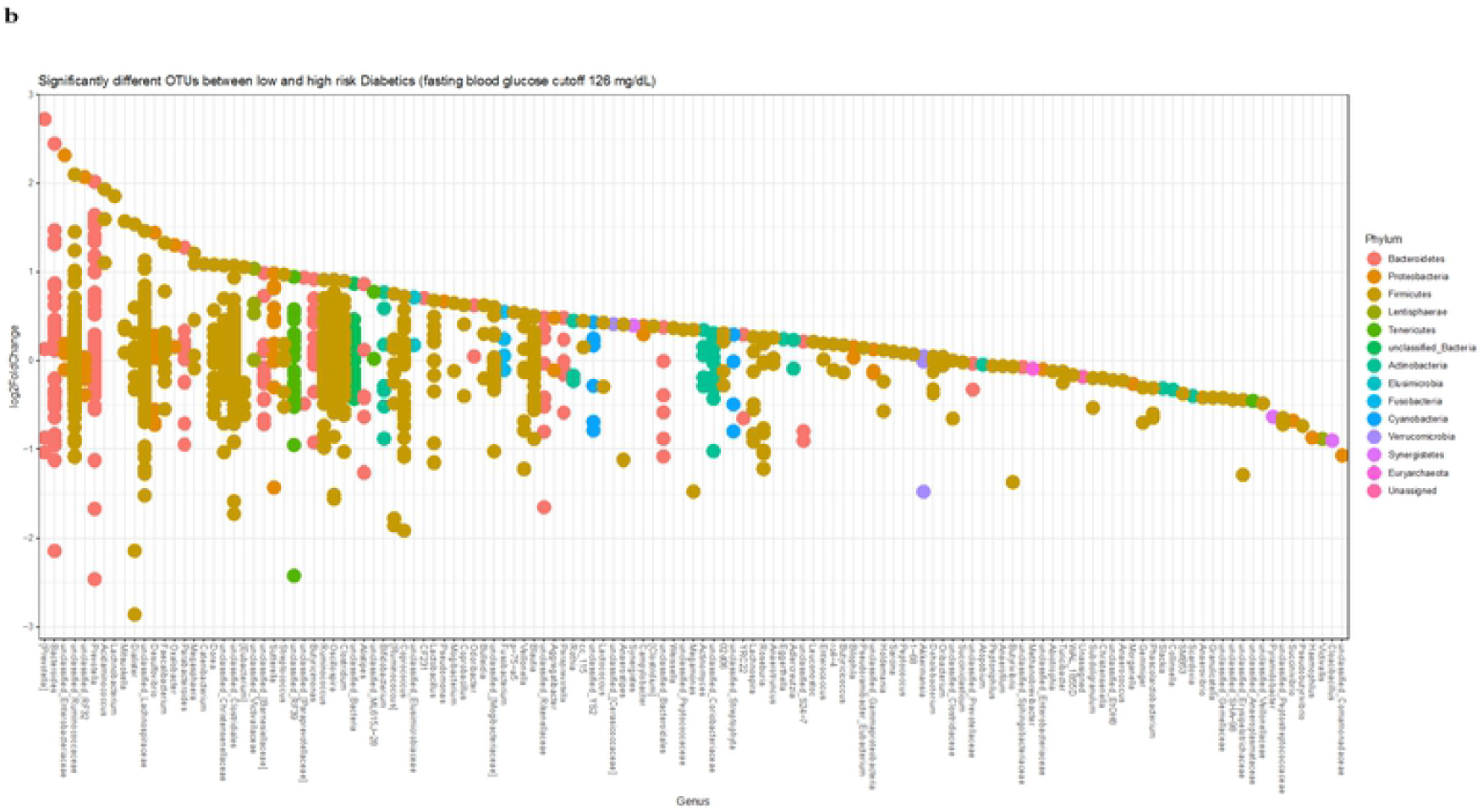
Fold Change plots of enriched OTUs for: T2D versus controls (a) and glucose levels for high versus low T2D status (b). An overall positive enrichment of microbiota genus/families for diabetics compared to healthy individuals and amongst diabetic participants was observed. Those with high glucose levels exhibited slightly more positive enrichment compared to those at lower risk of fasting hyperglycemia.

## Discussion

In this study we performed the largest microbiome study ever conducted in Saudi Arabia, as well as the first-ever characterization of gut microbiota T2D versus non-T2D in this population. We used shotgun metagenomic sequencing to obtain 16S rRNA reads identifiable down to genus level from the stool samples of 461 T2D and 119 non-T2D Saudi participants from the Eastern Province of Saudi Arabia, a region particularly affected by T2D [15]. We assessed the microbiota abundance based on diabetes status and glucose levels, and examined community diversity patterns to compare with other T2D microbiota studies from around the globe. These efforts are important and warranted given the scarcity of microbiome data in Middle Eastern populations, and these results provide a useful addition to the global microbiome reference dataset in an under-examined community. Saudi Arabian T2D costs have risen over 500% in two decades with 10 million individuals estimated to be diabetic or pre-diabetic, therefore comparing overlapping and varying patterns in gut microbiota with other studies is critical to assessing novel treatment options in light of a rapidly growing T2D health epidemic [15-16].

Community level differences are evident in the Saudi population between T2D and non-T2D individuals, and diversity patterns appear to vary from well-characterized microbiota from Western cohorts. Indeed, in contrast to Western cohorts that often show associations between decreased gut microbiota diversity and insulin resistance, here we show that Saudi participants with T2D exhibited higher relative diversity in comparison to normal metabolic counterparts [17]. These results are similar to a recent report from Al Bataineh and colleagues who characterized microbiomes in a cohort of 50 T2D and non-T2D individuals from the United Arab Emirates, though higher diversity in that smaller T2D cohort was determined to be insignificant when controlling for age [14]. Sex was not found to play a role in community structural differences, and results were independently validated between females and males. The role of overall community diversity decreasing in T2D populations has been widely cited in early studies on Western populations, yet larger meta-analyses involving global populations have distorted this pattern and highlight the importance of locally representative studies [18].

We observe significant differences between T2D and non-T2D individuals for many microbial taxa, as well as between T2D individuals with high and low fasting blood glucose levels. Concordant with studies conducted on Western populations is the association of increasing Bacteroidetes/Firmicutes ratio with T2D and in our overweight and obese T2D cohort, increased Bacteroidetes may be functionally related to metabolism of branched chain amino acids which has been linked to obesity-related metabolic phenotypes [3, 19]. Among OTUs assigned at the genus taxonomic level, *Prevotella* and *Bacteroides* OTUs showed some of the most significant log-fold increases in abundance for diabetics (over four-fold increases in abundance), species of which have been functionally associated with the development of insulin resistance and glucose intolerance [20]. Among Firmicutes however, levels of *Acidaminococcus* and *Megasphaera* were positively correlated with T2D, as has been previously observed, and could functionally relate with increases to Bacteroidetes through complementary amino acid metabolism [21-22]. We observed higher levels of *Akkermansia* in the Saudi T2D group, despite potential protective effects for obesity and metabolic disease. Associations of levels of *Akkermansia*, a mucus-consuming taxon, have been observed to be associated with health and with ethnicity in Western populations and may represent an impact of dietary and lifestyle effects on microbiota composition, as this microbe is rarely observed in more traditional cultures across large geographic regions [23]. It should be noted however that *Akkermansia* levels are also often increased in response to metformin intake in T2D subjects (metformin use metadata is not known for the current cohort) [24]. Taxonomic differences associated with T2D likely reflect shared or complementary functional and metabolic traits but may be regionally specific based on dietary and environmental variations known to influence the microbiome [23-25].

Based on diabetes status and quantified glucose levels of Middle Eastern participants, relatively stable differences in stool composition were observed by differential abundance and alpha diversity measures. Many studies have examined T2D associations with gut microbiota in populations around the globe, and while some patterns generally validate across studies such as individual taxon abundance variation, others such as overall community diversity do not replicate consistently. Obesity, diet, lifestyle and ancestry are all factors that influence T2D and each varies significantly from culture to culture around the globe, meaning that the patterns in T2D development and roles of the microbiome likely vary as well. As a rapidly emerging chronic condition in Saudi Arabia and the Middle East, T2D burdens have grown more quickly and affect larger proportions of the population than any other global region, making a regional reference T2D-microbiome dataset critical to understanding the nuances of disease development on a global scale.

## Materials and Methods

### Study Populations

Between 2015-2019, stool samples and data were collected from 461 consecutive diabetic patients attending the Diabetic Clinics, King Fahd Hospital of the University, Al-Khobar, Saudi Arabia and from 119 healthy controls. Participants ranged in age from 30-75 years and had a body mass index (BMI) ranging from 27 to 40 kg/m^2^. The T2D patients had a minimum disease duration of 5 years. Table 1 outlines the patient demographics and clinical characteristics. Baseline measurements included anthropometric measurements, physical examinations and in-person surveys. Participants who had been treated with antibiotics in the previous three months, were pregnant or lactating, or had inflammatory bowel disease were excluded from the study. Blood and stool samples were collected from participants and were stored immediately after collection at −80 °C. Ethical approval of the study was obtained from the local Institutional Review Board (IRB) committee and the study was conducted according to the ethical principles of the Declaration of Helsinki and Good Clinical Practice guidelines (IRB-2019-01-112). All participants provided written informed consent.

**Table 1.** Clinical and demographic characteristics for Saudi Arabian T2D cases (n-461) and controls (n=119).

|  | **Ratio** | **Male** | **Female** |
| --- | --- | --- | --- |
| **Gender** | 1: 0.83 | 54.50% | 45.50% |
|  | **Mean ± SD** |  |  |
|  | **Total** | **Male** | **Female** |
| **Age (Years)** | 52.6±8.83 | 51.82±9.28 | 53.5±8.25 |
| **Glucose(mg/dl)** | 165.7±68.89 | 161.45±57.71 | 166.8±74.09 |
| **HBA1c (%)** | 8.55±1.76 | 8.45±1.65 | 8.65±1.85 |
| **Duration (Years)** | 3-25 | 4-25 | 3-22 |
| **BMI (kg/m<sup>2</sup>)** | 27-40 | 27-37 | 30-40 |

### Methods for DNA library preparation and sequencing

Sample collection and microbial DNA extraction were standardized to minimize confounding effects of the technical procedure. Stool samples were taken from T2D (n=461) and from healthy (n=119) participants. Fecal samples were provided by the patients whilst attending the outpatient clinic and immediately stored at −20°C. The samples were subsequently transported on dry ice to the research laboratory where they were stored at −80°C. Bacterial DNA extraction from stool samples was performed using QIAamp Fast DNA Stool Mini Kit (Qiagen, Hilden, Germany) according to the manufacturer’s instructions. In brief, approximately 200 mg of stool was placed in a 2 ml microcentrifuge tube and kept on ice. InhibitEX Buffer (1 ml) was added to each stool sample, homogenized thoroughly by vortexing and incubated at 70°C for 5 minutes. Each sample was centrifuged (20,000 g) for minute and 200 µl supernatant was pipetted into 1.5 ml microcentrifuge tube containing 15 µl proteinase K and 200 µl lysis buffer and incubated at 70°C for 10 minutes. This was followed by the addition of 200 µl of ethanol and mixed by vortexing. The lysate (600 µl) was transferred to the QIAamp spin column and centrifuged (20,000 g) for 1 minute. Finally, the QIAamp spin column was opened and washed twice with two different washing buffers. The DNA was eluted into a new 1.5 ml microcentrifuge tube by adding 200 µl elution buffer. DNA samples were checked for purity using the Nanodrop 2000 Spectrophotometer (ThermoFisher Scientific). Three independent extractions were performed from each sample to ensure robust representation of all microbial content. DNA was stored at −80 °C till the time of processing.

Sequencing was performed using either the Swift Amplicon 16S panel (Swift Biosciences) or a custom protocol. For the Swift protocol, 20 ng of stool-derived DNA was used for 16S sequencing library preparation using the 16S Primer Panel v2, the Swift Normalase Amplicon Panels (SNAP) Core Kit, and the SNAP Combinatorial Dual Index Primer Kit (Sets 1A and 1B) (Swift Biosciences, CA). The indexed libraries were on average 620 base pairs (bp) in length, and individual DNA libraries were diluted to 2.5 nM, pooled in equimolar proportion, and sequenced on a NovaSeq 6000 SP flow cell (Illumina, CA) using 250 bp paired-end reads. For the custom approach, PCR was performed on each sample using the 515F primer (forward primer) and one of the 100 806rcbc primers (reverse primer). These primers contained: sequence homologous to region V4 of the 16S rRNA in forward and reverse; Illumina adaptors; and the reverse primers contained indexing sequences. Taq PCR Master Mix from Qiagen was used to prepare the PCR master mix. A PCR reaction was performed on each extracted DNA sample, i.e. each stool sample had three PCR reactions. The PCR product was run on 1% agarose gel. The band of expected size (381bp) was excised from gel and purified with gel purification kit from Qiagen. The three PCR products from each sample were pooled together. The pooled and purified PCR product was quantified with NanoDrop 2000 (Thermo Sciences, USA).

Equal concentrations of DNA from each sample (5ng of DNA) were pooled together. For each sequencing run, DNA from 50 samples was pooled to make the DNA library for each batch. The final concentration of the DNA library was quantified with real time PCR using the Kapa library quantification kit (Roche, USA) according to the manufacturer’s instructions. The DNA library of each batch was sequenced using the MiSeq platform from Illumina (Illumina, USA) using the MiSeq reagent V2 500cycles Kit from Illumina and the custom read1 (TATGGTAATTGTGTGCCAGCMGCCGCGGTAA), read2 (AGTCAGTCAGCCGGACTACH VGGGTWTCTAAT) and index (ATTAGAWACCCBDGTAGTCCGGCTGACTGACT) sequencing primers. PhiX DNA (Illumina, USA) was used as a control library.

### Analyses

Figure S1 overviews the analytical pipeline and workflow employed for these analyses. 16S rRNA (V4 region) sequences were used in this study and sequenced with Illumina software which handled the initial primer and barcode processing of all raw sequences. Raw sequences were demultiplexed with Illumina’s bcl2fastq2 v2.20 [26]. FastQC was then used for further processing to remove samples with low quality scores across the majority of bases [27]. After de-multiplexing the raw sequences and screening via FastQC, the majority of data processing was executed in QIIME2 with custom scripts. Paired-end reads were joined using VSEARCH. Chimera amplicon removal and abundance filtering were processed using Deblur [28]. Amplicon sequences were clustered and assembled into Operational Taxonomical Units (OTUs) using closed reference clustering against the Greengenes 13_8 database via VESEARCH. Taxonomic assignment was performed using a pre-trained Naïve Bayes classifier with Greengenes OTU database. The abundance tables and data obtained from QIIME2 were combined into a Phyloseq object and further analyzed in R with custom scripts [29].

## Acknowledgement

The authors would like to acknowledge the financial supported extended by King Abdulaziz City for Science and Technology, Riyadh, Saudi Arabia, grant numbers 12-MED2799-46 and 13-MED1881-46. We are also grateful to the nurses and technical staff for their work and dedication.

## Supporting information

**Fig S1:** Data processing and analyses pipeline for Saudi T2D 16S microbiota study.

**Fig S2:** Principal coordinate analyses of 16S microbiota data from Saudi T2D and control participants using: (a) sex and (b) T2D status.

**Fig S3a:** Heatmap of top 150 genus for (a) non-T2D and (b) T2D (OTU abundance based on BrayCurtis dissimilarity).

**Fig S3b:** Heatmap of top 50 genus for (a) non-T2D and (b) T2D individuals listed, respectively.

**Fig S4a:** Heatmap of top 150 gut microbiota 16S genus for (a) T2D and (b) T2D in Saudi females (OTU abundance based on BrayCurtis dissimilarity.

**Fig S4b**: Abundance of gut microbiota 16s taxonomic composition of: a) non-T2D vs (b) T2D in Saudi females

**Fig S5a:** Heatmap of top 150 gut microbiota 16S genus: (a) non-T2D (b) T2D in Saudi males (OTU abundance based on BrayCurtis dissimilarity).

**Fig S5b:** Abundance of gut microbiota 16S taxonomic composition of: (a) non-T2D versus (b) T2D in Saudi males.

**Fig S6:** Heatmap of top 150 genus for Saudi 16S gut microbiota for individuals with: (a) < 126 mg/dL and (b) >126 mg/dL (OTU abundance based on BrayCurtis dissimilarity).

**Fig S7:** Alpha diversity 16S gut microbiota assessment in Saudi males and females using: Chao1, ACE, Shannon-Weaver, Simpson, Inverse Simpson and Fisher indices.

**Fig S8:** Shannon and Simpson Alpha diversity: (a) T2D versus (b) non-T2D status.

**Fig S9:** *Bacteroidetes-Firmicutes* ratio in Saudi non-T2D cases and controls using 16S gut microbiota data.

**Table S1:** The most divergent microbiota genus between Saudi T2D cases and controls (a) and between T2D cases with high (> 126 mg/ dL) and low (< 126 mg/ dL) glucose (b). Positive 16S fold change indicates upregulation in diabetics.

